# How to Use Gravity to Accelerate Bone Adaptation: A Computational/Experimental Investigation of Exercises for Bone

**DOI:** 10.1101/2024.05.10.593555

**Authors:** Andrew R. Wilzman, Devin Tomoko Wong, Karen L. Troy

## Abstract

Impact exercises are known to increase bone mineral density (BMD) and in turn, bone strength and resistance to fracture. The biochemical pathways driving changes in BMD take months to complete, complicating our ability to understand how specific exercises influence the remodeling stimulus received by the bone. The purpose of this study was to compare several measures that have been theoretically linked to bone remodeling stimulus, including accelerations measured by Inertial Measurement Units (IMUs) at the middle of the tibia, ground reaction forces measured by force plate, joint contact forces estimated by musculoskeletal modeling, and tibia strains estimated by finite element modeling informed by high-resolution CT imaging. Twenty healthy adults (10 male: 22.1 +/- 2.2 years; 10 female: 21.3 +/- 1.3 years) participated in a biomechanical investigation of how drop height and landing style (bilateral vs. unilateral) affect the various bone remodeling stimuli. The results showed that while drop height consistently had significant direct relationships with stimulus magnitude, there was little benefit to drop heights greater than 0.4 m. In contrast, switching from a bilateral to a unilateral landing had a large positive effect. The stimuli calculated based on IMU data showed opposite trends compared to force plate and musculoskeletal modeling-based calculations, highlighting the need for caution in how IMUs are placed, and the resulting data interpreted, in the context of bone loading. A post-hoc analysis showed that a linear regression with predictor variables of kinematics, jump height, landing type (unilateral vs. bilateral) and the Ground Reaction Force FFT Integral could explain 79% of the variance in the bone remodeling stimulus that was predicted using much more sophisticated (and labor intensive) modeling. We conclude that higher level biomechanical modeling may not be necessary to understand the magnitude of a bone remodeling stimulus of an exercise.

## Introduction

To load, but not overload; to use, but not overuse. These are the recommendations we offer those who seek to improve their skeletal strength, not just for a reduced risk of fracture now, but later. Osteoporosis is a skeletal disorder defined by a reduction in bone mineral density (BMD) due to old age. This systemic BMD loss results in weak skeletal tissue, and an overall increased risk of bone fracture^1^. Osteoporosis affects women at a higher rate than men, with approximately 20% of women and 12% of men across the world^2^. A recent large-scale review of Medicare spending data in the United States found a total increase of over $30,000 in spending for patients with bone fractures due to osteoporosis in the first year after fracture^3^. Osteoporosis is currently diagnosed by routine Dual-energy X-ray Absorptiometry (DXA) scans that measure bone mineral density in various parts of the body^4^, and treated with medications such as bisphosphonates^5^. The prevalence of osteoporosis places a significant burden on healthcare spending, but the real impact of the disease is a continuously decreasing capability to maintain independence and quality of life in the aging population.

Osteoporosis is most effectively prevented by optimizing peak bone mass, which occurs during young adulthood^6^. Bone tissue is known to adapt to mechanical loading through an adaption mechanism driven by osteocytes that coordinate bone apposition, evidently targeting areas of high strain^7^. First proposed by Harold Frost in 1987, the “mechanostat” theory states that bone works to stiffen these areas of high strain to result in a structure optimized for the forces regularly imposed on the skeleton^8^. In a 2003 update, Frost wrote about bone’s ability to sense these forces as a signal and prioritize them based on frequency^9^. Since then, researchers have been eager to study and test the theory by measuring the osteogenic effects of new skeletal loads on bone structure and density, such as introducing new loading by adding jumping to exercise routines^10,11^.

It is widely accepted that bone adaptive response is diminished as the number of load repetitions increases^12^. Although theoretical models for this relationship have been tested and published, the specific input and parameter values needed to predict changes in bone strength given a particular skeletal exercise regimen are unknown^13–17^. In 2010, Ahola et al. devised and tested a Daily Impact Score (DIS) as an accelerometer-based remodeling stimulus^14^. Since then, more computationally intense methods have been developed and tested, particularly using Finite Element (FE) models^18–22^. While models have grown in complexity, it is unclear whether the additional cost of data and computation are justified by a different result. The key input value can be thought of as a theoretical bone remodeling stimulus, with the understanding that the stimulus decays with each repetition yet is cumulative over the time it takes the bone to adapt.

In this study, we focused on how to quantify the magnitude of the stimulus, and how a person could voluntarily manipulate this stimulus by modifying their activity or biomechanics. We also investigated how a theoretical bone remodeling stimulus would change, based on the amount and sophistication of the data available. We considered three levels of detail for the applied forces to the skeleton. The simplest measure is based on tibia acceleration, which can be easily captured with low-cost Inertial Measurement Units (IMUs). The next is vertical ground reaction force, whose measurement requires calibrated force platforms or pressure-sensing insoles. The most detailed force metric is an estimate of total joint contact force. This requires three-dimensional motion capture and musculoskeletal modeling to determine and sum the ground reaction force and estimated muscle force vectors. This force could then be applied to an FE model representing the tibia to calculate the bone’s strain reaction. The high cost of computation, expert labor, and high-precision measurements across multiple, expensive systems brought us to our final question: can the most accurate metric be predicted with less expensive inputs?

To answer the proposed questions, we designed an example bone loading exercise that would optimize bone impact with respect to energy required to complete a task set.

Dropping from elevated surfaces was selected as the task, taking advantage of gravity and simplicity. Thus, this study had two goals: (1) to determine the degree to which different landing methods could manipulate theoretical bone remodeling stimulus magnitudes, and (2) to understand the value of added complexity in estimating these stimuli. Our hypotheses for the first goal were that unilateral jump landings would produce higher stimulus magnitudes than bilateral landings and that jump height would be directly related to stimuli. Each participant will adopt their own landing strategy through sensitive, subconscious motor control feedback mechanisms. In these experiments, we will test if participants modulate the metrics that we associate with bone adaptation. For the second goal we hypothesized that a combination of less detailed inputs can be modeled to predict more detailed stimulus magnitudes. The only measurement tool studied here that is practical for use outside of controlled laboratory settings is the IMU that measures accelerations and rotational velocities. In this study, we only consider the DIS developed by Ahola et al. in 2010^14^. Since then, more sophisticated methods have been studied and developed to process IMU data for biomechanics research. While this is a limitation, we chose to narrow our study’s scope to determine if the IMU-fueled outcomes agree with more sophisticated methods, and more importantly if those outcomes could be used to determine the more sophisticated ones without having to directly measure them.

## Methods

### Data Collection

Twenty healthy adults (Table 1) gave written informed consent to participate in this institutionally approved study. Upon consent and meeting inclusion criteria for the study, we recorded their age, height, weight, sex, and musculoskeletal injury history. High resolution peripheral quantitative CT (HRpQCT, XtremeCT I, Scanco, Switzerland) was used to image a standardized 9.01 mm 3D region of each participant’s right distal tibia located 22.5 mm proximal to the distal subchondral plate (82 µm voxel size, 59.4 kV and 900 mA, effective radiation dose: 0.3 mSv).

**Table 1.** Participant demographics reported as mean (*standard deviation*)

|  | Male (n = 10) | Female (n = 10) |
| --- | --- | --- |
| Height (m) | 1.784 (0.043) | 1.628 (0.051) |
| Mass (kg) | 73.5 (7.5) | 55.4 (5.7) |
| Age (years) | 22.1 (2.2) | 21.3 (1.3) |

For the experiment, each participant removed their shoes and had at least five minutes to stretch and warm up before beginning motion tasks. Forty-one passive reflective markers were applied to the body using a modified plug-in gait marker set with extra clusters on the thighs and shanks. We measured skeletal segment trajectories using a ten-camera motion capture system (100 Hz, Vicon Motion Systems Ltd, UK), reaction forces with two six-axis force plates (1000 Hz, AMTI, Watertown, MA), and tibial acceleration with a dual IMU/EMG system adhered to the tibialis anterior (1000 Hz, Delsys Trigno Avanti, Delsys Inc., Natick, MA). The tasks were completed in the order shown in Table 2.

**Table 2.** Participant task set.

| Task | Sets x Trials | Instruction |
| --- | --- | --- |
| Bilateral Drop | 3 x 3 | Jump down from 0.4, 0.5, and 0.6 m platforms onto force plates with both feet. |
| Unilateral Drop | 3 x 2 | Jump down from 0.4, 0.5, and 0.6 m platforms, landing on only the right foot. |

### Data Processing

The outcomes were designed to represent a stimulus magnitude from each participant during the measured task. They were divided into four sections, with increasing degrees of complexity of data and/or invasiveness to the person to collect. To model the contributions of both loading magnitude and rate^17,23^, for each outcome listed above we included the effects of loading or strain rate using the product of magnitude and rate^21^, or an integral of the timeseries’ Fast Fourier Transform (FFT) from 0 Hz up to the frequency that encompasses 90% of the total signal power^17,21^. The total pool of outcomes is detailed in Table 3.

**Table 3.**
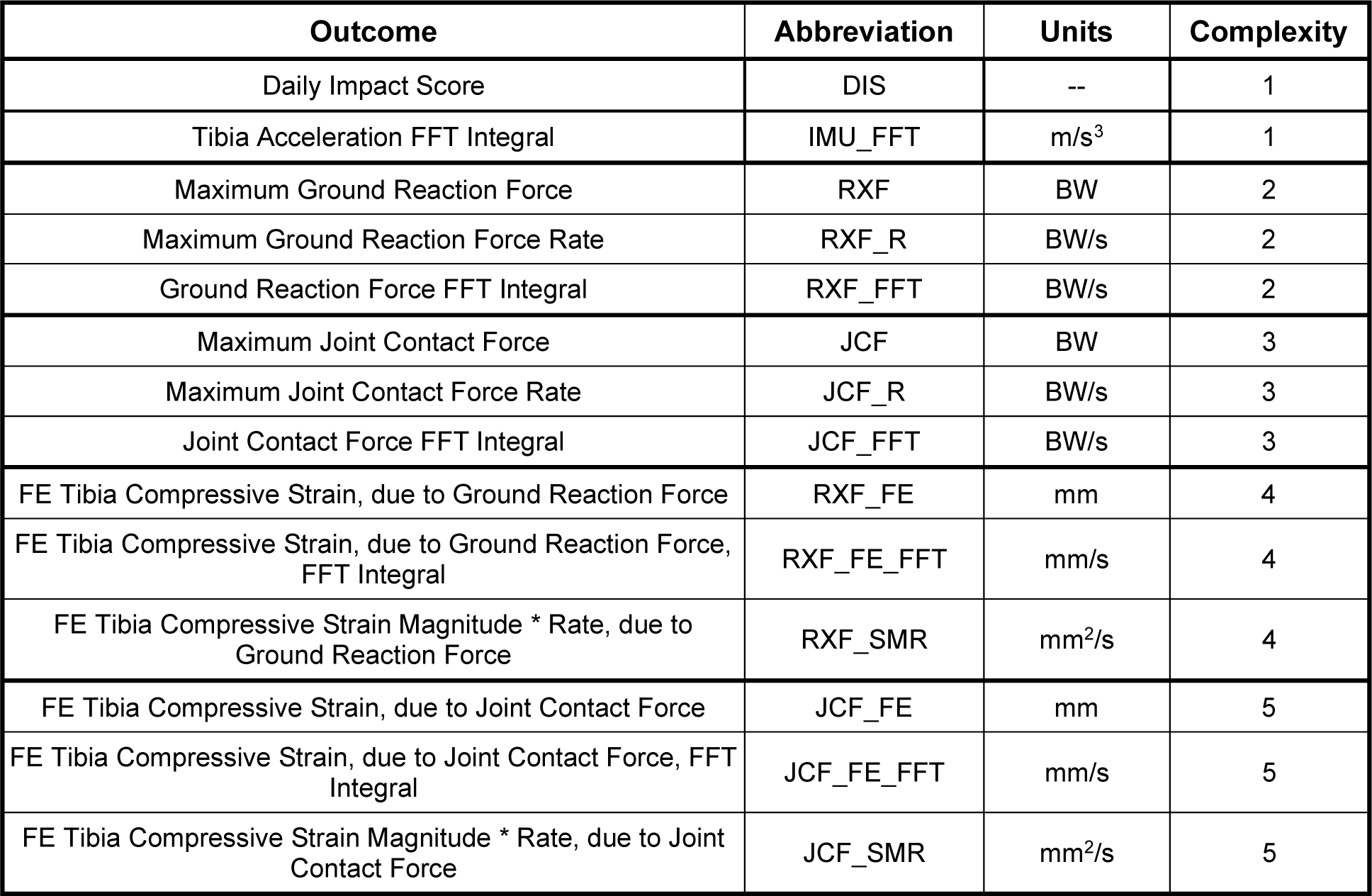
Outcome variables tested here, ranked by complexity.

| Outcome | Abbreviation | Units | Complexity |
| --- | --- | --- | --- |
| Daily Impact Score | DIS | -- | 1 |
| Tibia Acceleration FFT Integral | IMU_FFT | m/s<sup>3</sup> | 1 |
| Maximum Ground Reaction Force | RXF | BW | 2 |
| Maximum Ground Reaction Force Rate | RXF_R | BW/s | 2 |
| Ground Reaction Force FFT Integral | RXF_FFT | BW/s | 2 |
| Maximum Joint Contact Force | JCF | BW | 3 |
| Maximum Joint Contact Force Rate | JCF_R | BW/s | 3 |
| Joint Contact Force FFT Integral | JCF_FFT | BW/s | 3 |
| FE Tibia Compressive Strain, due to Ground Reaction Force | RXF_FE | mm | 4 |
| FE Tibia Compressive Strain, due to Ground Reaction Force, FFT Integral | RXF_FE_FFT | mm/s | 4 |
| FE Tibia Compressive Strain Magnitude * Rate, due to Ground Reaction Force | RXF_SMR | mm<sup>2</sup>/s | 4 |
| FE Tibia Compressive Strain, due to Joint Contact Force | JCF_FE | mm | 5 |
| FE Tibia Compressive Strain, due to Joint Contact Force, FFT Integral | JCF_FE_FFT | mm/s | 5 |
| FE Tibia Compressive Strain Magnitude * Rate, due to Joint Contact Force | JCF_SMR | mm<sup>2</sup>/s | 5 |

#### Tibia Acceleration

A Delsys Trigno Avanti sensor affixed to the right anterior shank was used to measure tibial acceleration (sampling rate: 1000 Hz), and the data were passed through a 50 Hz, fourth-order low-pass Butterworth filter. The maximum acceleration magnitude was binned according to the Daily Impact Score (DIS)^14^. The integral of the Fourier transform was also recorded (IMU FFT).

#### Ground Reaction Force

Ground reaction force was measured using a 1000 Hz, 6-axis force plate embedded into the landing surface. The data were passed through a 300 Hz fourth-order low-pass Butterworth filter before recording maximum reaction force (RXF), maximum force rate (RXF_R), and the integral of the Fourier transform (RXF_FFT).

#### Joint Contact Force

The motion capture data were filtered with a 6 Hz fourth-order low-pass Butterworth filter before being exported to Visual 3D (C-Motion, Gaithersburg, MD) to model body segments and calculate inverse kinematics and dynamics. We filtered the data again and exported the result to OpenSim 4.4^24,25^, using model Gait 2392 as the basis for our calculations^26^. After model scaling, we used static optimization to estimate lower extremity muscle forces for each trial. Finally, the result was filtered again and total joint contact force at the ankle was calculated by summing the forces of all muscles crossing the ankle joint along with the net joint force calculated from inverse dynamics. The maximum ankle joint contact force (JCF), maximum force application rate (JCF_R), and integral of the Fourier transform (JCF_FFT) were recorded.

#### Tibia Compressive Strain

Subject-specific FE models of a 9.01 mm section of the distal tibia were created from HRpQCT images. A Scanco built-in FE solver was used with a single tissue model^27^ to estimate axial tibia stiffness (in N/mm) under platen compression. Reaction strain (RXF_FE) and contact strain (JCF_FE) were calculated by dividing RXF and JCF, respectively, by axial tibia stiffness. The time series were also analyzed to extract maximum tibial strain rates^21^. This produced the last outcomes: reaction and contact strain magnitude*rates (SMR) and integral Fourier transforms (RXF_SMR, JCF_SMR, RXF_FE_FFT,JCF_FE_FFT).

### Statistical Analysis

We initially calculated Pearson correlations for each outcome variable to assess their relationships. We expected the outcomes to be highly correlated within each complexity group; however, the correlations outside of these groups shed light on how the different means of measurement produce different estimates of a bone adaptation stimulus. We also analyzed correlations between key kinematic predictor variables: joint angle at contact and joint range of motion for the hip, knee, and ankle of the analyzed leg.

Our first hypothesis was that unilateral jump landings would produce higher stimulus magnitudes than bilateral landings. To test this, we used a paired t-test to compare each stimulus metric between landing conditions. We set our alpha significance threshold to 0.05 as a standard for this and the remaining statistical tests and corrected for multiple comparisons using Bonferroni correction.

Our second hypothesis was that jump height would be directly related to stimulus magnitudes. To test this, we regressed each outcome with jump height as a predictor, adding the number of landing limbs as a secondary predictor if it was found to be significant. Significance was assessed by testing the coefficient of determination (R^2^) with an F-statistic and each individual predictor with a t-statistic.

Our third hypothesis was that a combination of less detailed inputs could be modeled to predict more detailed stimulus magnitudes. To test this, we employed the Least Absolute Shrinkage and Selection Operator (LASSO) to choose predictor variables in the regression for each outcome. The pool of possible predictors was restricted to variables with lower complexity, including the kinematic predictors as a baseline. If height and number of landing legs were deemed significant, these were added as predictors. Data were split into training (80%) and testing (20%) sets for validation. During training, the data were normalized to a z-score as a preprocessing step and each feature’s scalar was saved to be applied on the predictors of the testing set. It is important to note that the scalar term is driven only by the training set data. Model performance was assessed on the test set by Mean Absolute Error (MAE) in units of standard deviations and R^2^.

## Results

### Overview

Pearson correlations of outcomes within the same complexity group were generally high and significant (Figure 1). The IMU-related outcomes showed strong negative correlations with the rest of the measures.

**Figure 1.**
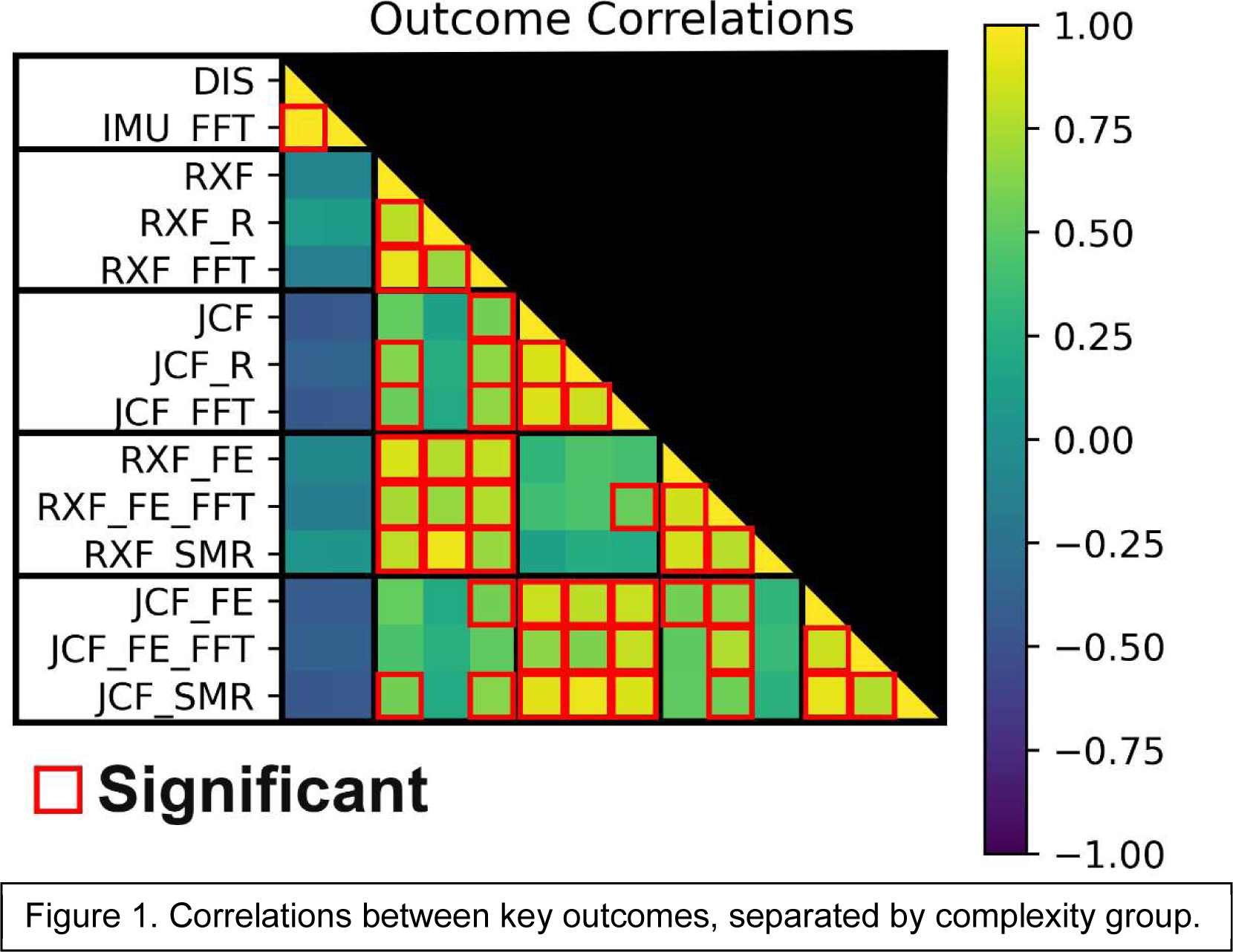
Correlations between key outcomes, separated by complexity group.

### Effect of unilateral vs. bilateral jumps on stimulus magnitude

In general, unilateral jump landings produced significantly higher stimulus magnitudes than bilateral landings. However, while the daily impact score and IMU FFT outcomes were significantly different between the landing groups, the relationship was reversed (Figure 2).

**Figure 2.**
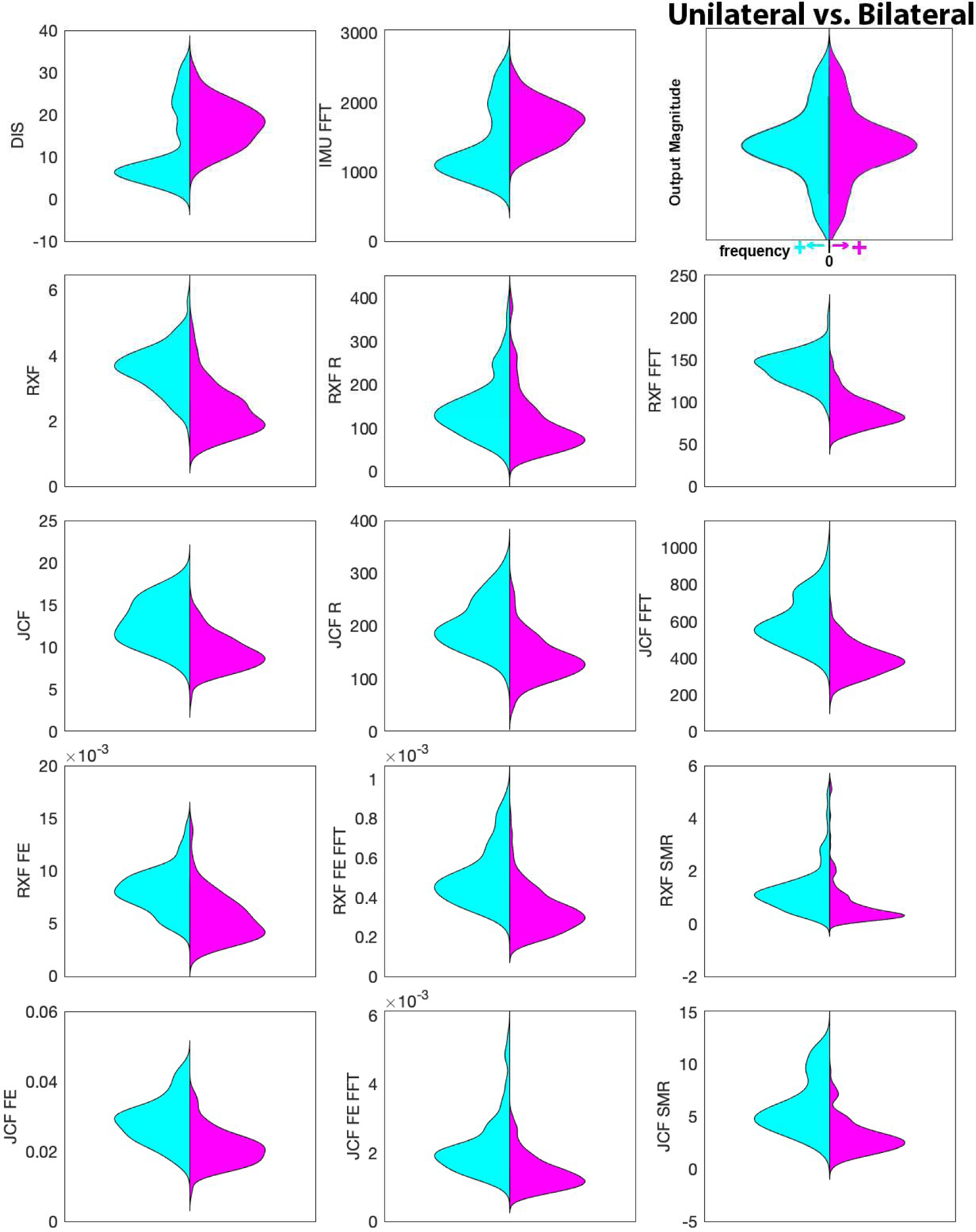
Comparison of unilateral and bilateral jump landings across all trials. Each graph shows a histogram that represents the frequency (x axes) of each output magnitude (y axes), mirrored across unilateral and bilateral landings. All comparisons were significant.

### Effect of jump height on stimulus magnitude

For most metrics, the 0.2 m height produced significantly lower outcomes than other heights. Fewer outcomes were significantly different in comparison with heights at 0.4 m, but 2 out of 3 outcomes in the highest complexity group (JCF_FE, JCF_SMR) were significantly higher in heights 0.6 m than 0.4 m. Upon further analysis, it was revealed that the differences in the unilateral trials drove this significance (Figure 3). To determine the relative influence of height vs. landing limbs on each outcome, post-hoc analysis included fitting each of them to a bivariate linear model with height and landing limbs as predictors. Both of predictors were significant. The details within the model accuracies demonstrate that the number of landing limbs explains more variance in each of the outcomes than height, apart from RXF_R (Table 4).

**Figure 3.**
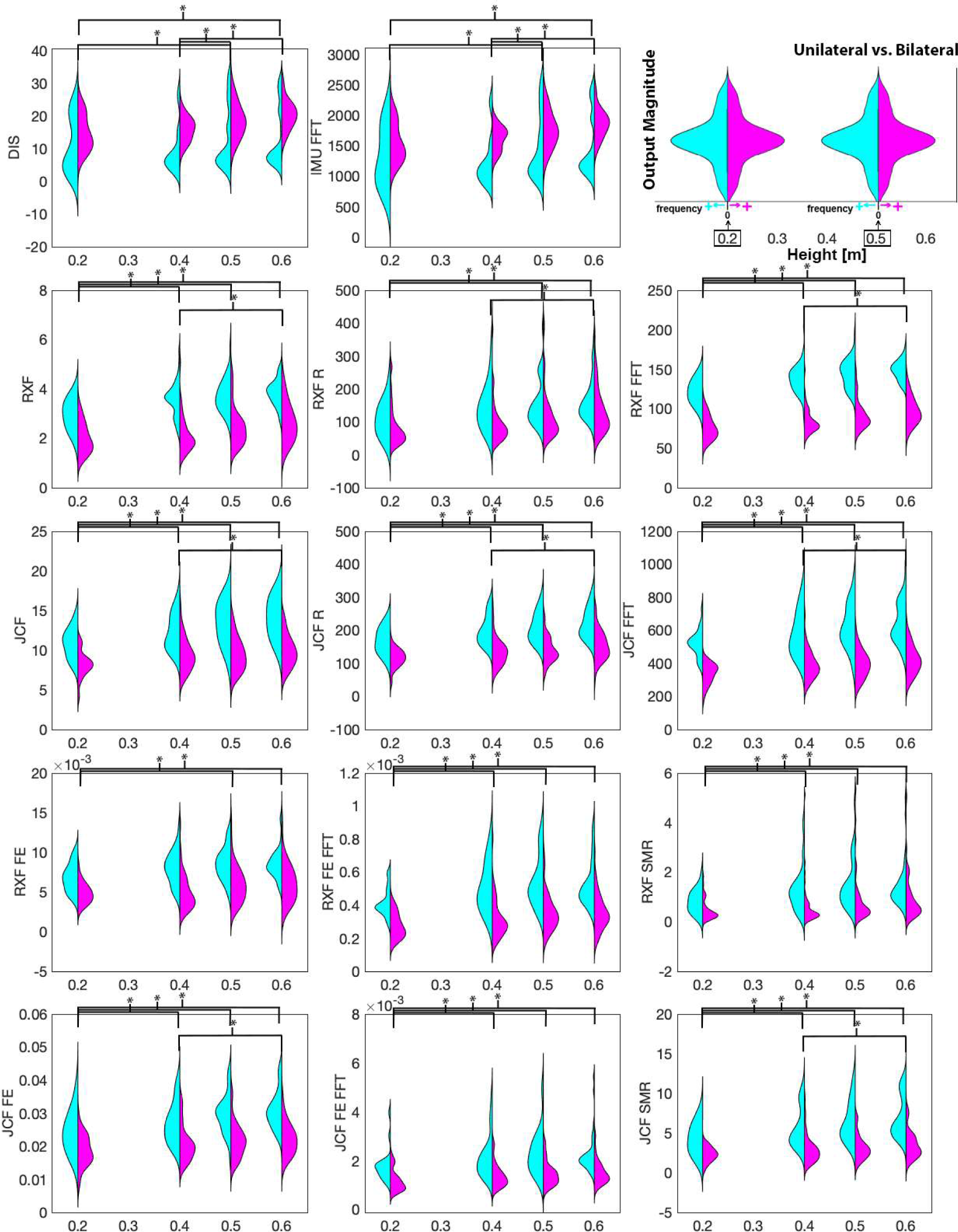
Comparison of unilateral and bilateral jump landings, separated by height. Each histogram represents the frequency of outputs, mirrored with respect to unilateral and bilateral landings. Significant comparisons are denoted above.

**Table 4.** Comparison of the relative influence of drop height and landing limbs on each outcome. All coefficients were significant with p < 0.05 and Bonferroni correction.

| Outcome | Complexity Group | Coefficients ($\beta$) | | Coefficients of Determination ($R^2$) | Total Model $R^2$ |
| --- | --- | --- | --- | --- | --- |
| DIS | 1 | Drop Height | 10.96 | 0.049 | 0.192 |
|  |  | Landing Limbs | 5.60 | 0.140 |  |
| IMU_FFT | 1 | Drop Height | 667.21 | 0.057 | 0.199 |
|  |  | Landing Limbs | 314.68 | 0.139 |  |
| RXF | 2 | Drop Height | 1.88 | 0.085 | 0.485 |
|  |  | Landing Limbs | -1.23 | 0.411 |  |
| RXF_R | 2 | Drop Height | 131.46 | 0.070 | 0.131 |
|  |  | Landing Limbs | -35.68 | 0.065 |  |
| RXF_FFT | 2 | Drop Height | 60.10 | 0.085 | 0.730 |
|  |  | Landing Limbs | -50.87 | 0.659 |  |
| JCF | 3 | Drop Height | 5.68 | 0.085 | 0.444 |
|  |  | Landing Limbs | -3.50 | 0.370 |  |
| JCF_R | 3 | Drop Height | 109.74 | 0.088 | 0.422 |
|  |  | Landing Limbs | -63.97 | 0.345 |  |
| JCF_FFT | 3 | Drop Height | 248.49 | 0.064 | 0.542 |
|  |  | Landing Limbs | -208.86 | 0.489 |  |
| RXF_FE | 4 | Drop Height | $3.81 \times 10^{-3}$ | 0.050 | 0.311 |
| | | Landing Limbs | $-2.57 \times 10^{-3}$ | 0.267 | |
| RXF_FE_FFT | 4 | Drop Height | $2.37 \times 10^{-4}$ | 0.052 | 0.369 |
| | | Landing Limbs | $-1.74 \times 10^{-4}$ | 0.324 | |
| RXF_SMR | 4 | Drop Height | 1.31 | 0.042 | 0.129 |
|  |  | Landing Limbs | -0.54 | 0.090 |  |
| JCF_FE | 5 | Drop Height | $1.24 \times 10^{-2}$ | 0.070 | 0.365 |
| | | Landing Limbs | $-7.46 \times 10^{-3}$ | 0.303 | |
| JCF_FE_FFT | 5 | Drop Height | $9.92 \times 10^{-4}$ | 0.038 | 0.269 |
| | | Landing Limbs | $-7.33 \times 10^{-4}$ | 0.237 | |
| JCF_SMR | 5 | Drop Height | 4.65 | 0.071 | 0.396 |
|  |  | Landing Limbs | -2.93 | 0.334 |  |

### Effect of input data complexity on stimulus magnitude

Because height and number of landing legs affected each outcome, these were included as predictors in the LASSO regression models. Our first set of models included only kinematic variables as predictors for each regression as a baseline model (Table 5). Next, for complexity groups 2 and higher, we included the result of the outcomes in all lower complexity groups in the predictor pool (Table 6).

**Table 5.** Testing results for LASSO regression on each outcome using kinematics, height, and landing legs.

| Outcome | Complexity | MAE ($\sigma$) | R<sup>2</sup> |
| --- | --- | --- | --- |
| DIS | 1 | 0.62 | 0.41 |
| IMU_FFT | 1 | 0.65 | 0.39 |
| RXF | 2 | 0.53 | 0.53 |
| RXF_R | 2 | 0.65 | 0.09 |
| RXF_FFT | 2 | 0.40 | 0.69 |
| JCF | 3 | 0.52 | 0.63 |
| JCF_R | 3 | 0.54 | 0.62 |
| JCF_FFT | 3 | 0.39 | 0.76 |
| RXF_FE | 4 | 0.56 | 0.34 |
| RXF_FE_FFT | 4 | 0.59 | 0.39 |
| RXF_SMR | 4 | 0.53 | 0.00 |
| JCF_FE | 5 | 0.50 | 0.53 |
| JCF_FE_FFT | 5 | 0.45 | 0.47 |
| JCF_SMR | 5 | 0.52 | 0.57 |

**Table 6.** Testing results for LASSO regression on each outcome using lower complexity outcomes, kinematics, height, and landing legs.

| Outcome | Complexity | MAE ($\sigma$) | R<sup>2</sup> |
| --- | --- | --- | --- |
| RXF | 2 | 0.51 | 0.52 |
| RXF_R | 2 | 0.60 | 0.24 |
| RXF_FFT | 2 | 0.37 | 0.79 |
| JCF | 3 | 0.30 | 0.75 |
| JCF_R | 3 | 0.36 | 0.67 |
| JCF_FFT | 3 | 0.32 | 0.83 |
| RXF_FE | 4 | 0.17 | 0.92 |
| RXF_FE_FFT | 4 | 0.23 | 0.88 |
| RXF_SMR | 4 | 0.12 | 0.94 |
| JCF_FE | 5 | 0.22 | 0.91 |
| JCF_FE_FFT | 5 | 0.22 | 0.94 |
| JCF_SMR | 5 | 0.15 | 0.95 |

In the models that considered height, landing legs, and kinematics only, the outcomes in complexity group 1 were not modeled well by the LASSO regression.

The models that predicted FE outcomes found much better success on testing sets (Figure 4). These models tended to assign large coefficients to joint reaction and contact forces compared to the rest of the possible predictor variables. To compare how these FE outcomes could be predicted without musculoskeletal modeling (required to determine joint contact forces), we removed joint contact force data from the pool of predictors and found that it reduced the accuracy by 10-30% (Table 7).

**Figure 4.**
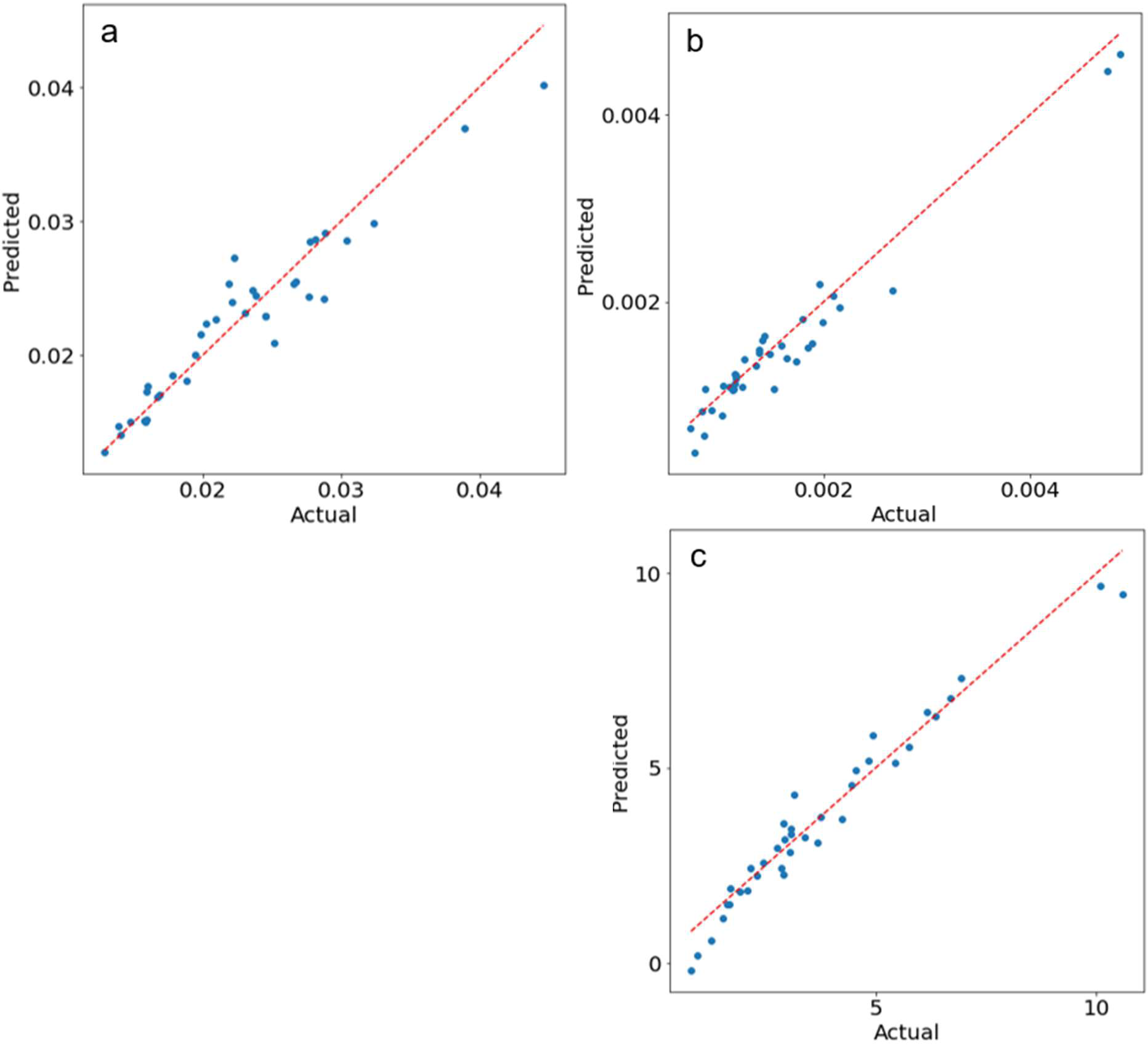
Actual Versus predicted joint Contact force variables on the testing A maximum MAE of 23% was observed among models trained to predict JCF_FE (a), JCP_FE_FFT (b), and JCF_SMR (c), For JCF_FE and JCF_FE_FFT, the LASSO retained much of its predictor pool, while JCP_SMR retained only the three Joint contact force variables, ankle ROM, hip flexion at contact, and DIS (Appendix A3)

**Table 7.** Testing results for LASSO regression on FE outcomes.

| Outcome | Without JCF data |  | With JCF data |  |
| --- | --- | --- | --- | --- |
| | MAE ($\sigma$) | R<sup>2</sup> | MAE ($\sigma$) | R<sup>2</sup> |
| RXF_FE | 0.34 | 0.83 | 0.17 | 0.92 |
| RXF_FE_FFT | 0.38 | 0.78 | 0.23 | 0.88 |
| RXF_SMR | 0.17 | 0.91 | 0.12 | 0.94 |
| JCF_FE | 0.40 | 0.70 | 0.22 | 0.91 |
| JCF_FE_FFT | 0.49 | 0.64 | 0.22 | 0.94 |
| JCF_SMR | 0.42 | 0.63 | 0.15 | 0.95 |

## Discussion

Here, we sought to understand the effects of jump height and number of landing limbs on the potential osteogenic stimulus. We found that landing on one leg, rather than increasing height, is a more effective way to generate a higher bone remodeling stimulus from drop landings. This agrees with an understanding that humans are more efficient at modulating impact loads bilaterally, while we meet unilateral tasks with higher joint stiffness to ensure stability^28^. While we considered altering the landing instruction set, we ultimately allowed each participant to control their landings as they deemed fit. However, one recent study found that a cue to land quietly can immediately reduce ground reaction force rates in runners^29^. Therefore, we recommend further studies to investigate how landing cues can manipulate these outcomes, particularly with the opposite approach of asking participants to land harder/louder.

The positive effect that jumping exercises have on bone mineral density are subtle, but clear in multiple intervention studies^30–32^. Utilizing the initial strength of participants, most intervention studies lean on healthy populations tasked with jumping to maximum heights or dropping from heights no higher than six inches. From the results of this computational analysis, we would expect to see higher increases in BMD in participants who drop from heights higher than 0.4 m. These findings are consistent with a 2018 review article that concluded ground reaction forces should exceed 3.5 * BW to be effective in increasing BMD through exercise intervention^32^. Across the board, our findings suggest that unilateral landings would be a more effective way to maximize increases in BMD (Figure 5).

**Figure 5.**
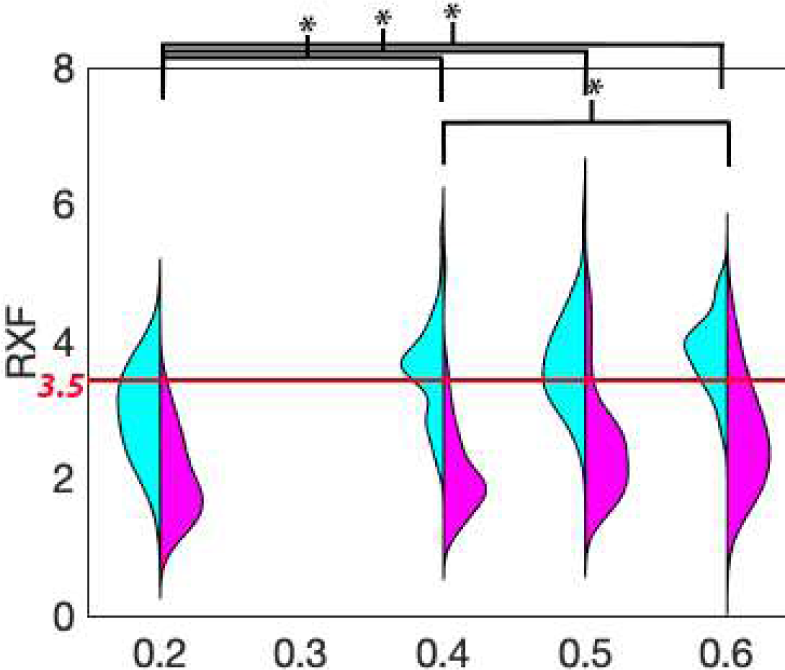
Reaction forces above 3.5 BW are seen in unilateral landings across the board, but only for heights above 0.4 m for bilateral landings.

The IMU data showed a clear opposition to the remainder of the outcomes, but still found its way into the chosen coefficients that predicted later outcome metrics. We speculate that this highlights a need for clear and robust fixation methods of IMU placement. Here, the IMU data were recorded on the same device as EMG data, which requires placement on a point of a muscle (the tibialis anterior) that maximizes cross-sectional area and minimizes skin occlusion distance. These devices were then subject to soft tissue artifacts not normally seen when applying devices that act wholly as IMUs, which are fixed to boney landmarks. The magnitude of these artifacts is almost certainly affected by the stiffness of the underlying tissues.

When the linear models were restricted to kinematic variables, 62-76% of variance in JCF could be predicted. However, when the same predictor pool was tasked to model FE outcomes it failed to reach an R^2^ greater than 60%. Interestingly, the mean error of the predictions remained within the same standard deviation range as their JCF counterparts. At minimum, ground reaction force data should be included to reliably predict the magnitude of an assumed osteogenic stimulus, highlighted by the success of unrestricted LASSO models tasked to model FE outcomes (Tables 6 and 7). On the other hand, RXF_FFT stood out as a metric that could be well modeled by kinematics and IMU data.

This analysis included several metrics that are not typically available outside research settings but that we assumed were good measures of osteogenic stimulus. We hypothesized that a combination of less detailed inputs could be modeled to predict more detailed stimulus metrics. The LASSO models explained 67% of the variance in joint contact force data if only kinematics and ground reaction forces are known. Once lower complexity outcomes are included as possible predictors, the model increased in accuracy with over 88% of the variance explained in estimated tibia strains. This could be explained by the inclusion of the forces that were used to estimate the tibia strain via FE estimated stiffness. Analysis of the coefficients chosen within these linear models (Appendix A) reveals a high degree of importance in using reaction forces and musculoskeletal modeling to predict the FE outcomes.

We expected that outcomes within the same complexity group would be highly correlated, and we attribute this correlation to measurements being derived from common means (by device and/or software). Correlations diminished outside of these groups, but the relationships that endured provided enough information to create linear models that predict FE outcomes without the need for CT imaging or any explicit FE modeling. This leads us to speculate that peak bone strains can be effectively estimated in musculoskeletal modeling software without FE solvers or bone density measurements. In other words, variance in bone strain was most closely associated with the applied loading, mainly because the present group of young adult participants had relatively homogenous tibia bone stiffness. Given the large changes in bone stiffness and architecture associated with aging and disease^33,34^, we believe it is still important to account for this factor when estimating osteogenic stimulus in other populations.

The first limitation of this study begins with the participant pool, which includes only healthy, mostly athletic, college students. Most of the participants were familiar with box jumps and landing strategies. A few participants were less likely to fully drop from the heights during unilateral tasks, using a partial “stepping” method instead of falling from the entire height of the box. However, this highlighted the real variation of task understanding when a person is asked to drop from a height onto one leg at a time. In the future, the “step down” version of this exercise could be completed using a handrail for balance, making it potentially suitable for many age and ability groups.

Another limitation of this study included the assumptions made to calculate the muscle force time series required to estimate joint contact forces. In OpenSim, static optimization was used for this, which may be less sophisticated than another tool, computed muscle control, that takes time dynamics into account^35^. We compared our results to While these may have resulted in slightly different results, the high accelerations of the drop landing caused too much noise in computed muscle control results to be reliable.

Finally, the FE outcomes were based on a standard, platen compressive finite element solver within Scanco’s XtremeCT software. The solver assumes that the material properties are linear-elastic and homogeneous and does not take bending stress into account^36,37^. These limitations may act together to decrease the validity of the strain metrics with respect to reality. However, the comparisons between the data remain valid through the standardization of protocol and use of subject-specific information.

## Conclusions

Unilateral drop landings significantly increase a theoretical bone remodeling stimulus compared to bilateral landings. Additionally, there may be little benefit to jumping from heights above 0.4 m when examining bone remodeling stimulus. However, outcomes reliant on IMU data showed the opposite effect and should be interpreted with caution. A reader should inquire about the placement of the IMU and whether soft tissue artifacts pose a risk to the integrity of the data.

Finally, within a homogenous group, bone remodeling stimulus can be accurately predicted through a combination of 3D motion capture, accurate ground reaction forces, and musculoskeletal modeling. While this is still a complex process, it may negate the need to use quantitative CT imaging and FE modeling to estimate a bone’s strain reaction due to a jump landing. The degree to which these theoretical remodeling stimulus calculations predict actual bone remodeling is yet to be prospectively tested in humans. These data and methods presented here make such future predictions possible.

## Acknowledgements

The authors would like to acknowledge Julia Nicolescu, Logan Gaudette, Elizabeth Bowman, and Madalyn Hague for their assistance in data collection and processing, Christopher Gens for his work in data processing, and Christopher Nycz for his support in motion capture data collection. The authors also acknowledge the Research Experiences for Undergraduates (REU) funding through the US National Science Foundation (NSF Award #2150076) for the opportunity to collaborate with undergraduate researcher Devin Wong.

## Appendix A. LASSO Regression

**Table A1.** LASSO regression model coefficients, bolded values are significant with p < 0.05 (Range Of Motion = ROM). Blank values represent removed variables.

| Outcome | Height | Landing Limbs | Hip Flexion ROM | Knee Flexion ROM | Ankle Flexion ROM | Hip Flexion at Contact | Knee Flexion at Contact | Ankle Flexion at Contact |
| --- | --- | --- | --- | --- | --- | --- | --- | --- |
| DIS | 1.89E+00 | 2.72E+00 | -1.78E+00 | 2.48E+00 | -2.66E+00 | 4.00E-01 | 3.91E+00 | 3.02E+00 |
| IMU_FFT | 1.14E+02 | 1.44E+02 | -1.06E+02 | 1.56E+02 | -1.57E+02 | 8.75E+00 | 2.07E+02 | 1.67E+02 |
| RXF | 3.58E-01 | -4.85E-01 | -1.69E-01 | -7.29E-02 | 4.86E-02 | -1.65E-01 | 8.84E-02 | 1.02E-01 |
| RXF_R | 2.07E+01 | -1.42E+01 | -9.64E+00 | 4.61E+00 |  | -1.02E+01 | 6.90E+00 |  |
| RXF_FFT | 9.39E+00 | -2.34E+01 | -2.95E+00 |  |  | -3.70E+00 |  | 1.83E-01 |
| JCF | 5.63E-01 | -1.70E+00 | -1.91E-02 | -7.51E-02 | 1.19E+00 | -1.43E-01 | -2.33E-01 |  |
| JCF_R | 1.67E+01 | -2.60E+01 |  | -1.19E+01 | 1.99E+01 | -8.22E+00 | -4.90E+00 | 6.41E-01 |
| JCF_FFT | 2.28E+01 | -1.07E+02 | -1.32E+01 | 1.62E+01 | 5.35E+01 |  | -1.84E+01 | -1.14E+01 |
| RXF_FE | 9.98E-04 | -7.26E-04 | -3.64E-04 | -8.48E-04 | 5.16E-04 | -1.91E-04 | 2.40E-04 | 3.80E-04 |
| RXF_FE_FFT | 3.35E-05 | -8.11E-05 |  | -8.87E-06 | 3.54E-05 | -1.36E-05 | 4.01E-06 |  |
| RXF_SMR | 2.95E-01 | -1.48E-01 | -1.80E-01 | -7.20E-02 | 7.72E-02 | -7.57E-02 | 1.64E-01 | 1.14E-01 |
| JCF_FE | 1.68E-03 | -3.02E-03 | 1.92E-04 | -2.18E-03 | 4.28E-03 | 1.41E-04 | -7.93E-04 |  |
| JCF_FE_FFT | 3.80E-05 | -4.13E-04 | 1.30E-04 | -7.58E-05 | 4.14E-04 | -2.85E-05 | -1.99E-04 | -1.70E-04 |
| JCF_SMR | 6.04E-01 | -1.27E+00 | 2.95E-01 | -7.85E-01 | 1.37E+00 | -2.48E-01 | -3.13E-01 |  |

**Table A2.** Testing results for LASSO regression models restricted to kinematic predictor variables.

| Outcome | Mean | Standard deviation ($\sigma$) | M.S.E. | M.A.E. | M.A.E. / $\sigma$ | $r^2$ |
| --- | --- | --- | --- | --- | --- | --- |
| DIS | 1.53E+01 | 7.25E+00 | 3.10E+01 | 4.50E+00 | 62.0% | 0.41 |
| IMU_FFT | 1.58E+03 | 4.08E+02 | 1.03E+05 | 2.64E+02 | 64.6% | 0.39 |
| RXF | 2.87E+00 | 9.61E-01 | 4.83E-01 | 5.09E-01 | 53.0% | 0.53 |
| RXF_R | 1.26E+02 | 7.12E+01 | 4.55E+03 | 4.60E+01 | 64.6% | 0.09 |
| RXF_FFT | 1.10E+02 | 3.13E+01 | 3.13E+02 | 1.25E+01 | 39.9% | 0.69 |
| JCF | 1.08E+01 | 2.89E+00 | 3.53E+00 | 1.51E+00 | 52.3% | 0.63 |
| JCF_R | 1.66E+02 | 5.47E+01 | 1.27E+03 | 2.94E+01 | 53.8% | 0.62 |
| JCF_FFT | 4.79E+02 | 1.49E+02 | 5.68E+03 | 5.82E+01 | 39.0% | 0.76 |
| RXF_FE | 6.65E-03 | 2.49E-03 | 3.04E-06 | 1.40E-03 | 56.1% | 0.34 |
| RXF_FE_FFT | 4.08E-04 | 1.53E-04 | 1.19E-08 | 9.00E-05 | 58.8% | 0.39 |
| RXF_SMR | 9.86E-01 | 9.12E-01 | 3.44E-01 | 4.84E-01 | 53.1% | -0.10 |
| JCF_FE | 2.43E-02 | 6.81E-03 | 1.86E-05 | 3.40E-03 | 50.0% | 0.53 |
| JCF_FE_FFT | 1.74E-03 | 7.54E-04 | 2.72E-07 | 3.43E-04 | 45.4% | 0.47 |
| JCF_SMR | 4.33E+00 | 2.55E+00 | 2.93E+00 | 1.32E+00 | 51.8% | 0.57 |

**Table A3-1.** LASSO regression model kinematic coefficients, bolded values are significant with p < 0.05 (Range Of Motion = ROM).

| Outcome | Height | Landing Limbs | Hip Flexion ROM | Knee Flexion ROM | Ankle Flexion ROM | Hip Flexion at Contact | Knee Flexion at Contact | Ankle Flexion at Contact |
| --- | --- | --- | --- | --- | --- | --- | --- | --- |
| RXF | 4.93E-01 | -4.49E-01 | -2.16E-01 | -2.14E-02 | 6.54E-02 | -1.44E-01 | 1.75E-01 | 2.16E-01 |
| RXF_R | 2.74E+01 | -1.32E+01 | -1.52E+01 | 9.69E+00 | 8.03E+00 | 3.10E-01 | 3.08E+01 | 1.90E+01 |
| RXF_FFT | 1.35E+01 | -2.27E+01 | -8.20E+00 | 3.89E+00 | 5.56E-01 | -2.08E+00 | 7.28E+00 | 6.83E+00 |
| JCF | 3.50E-01 | -1.67E-01 | -2.66E-01 | 5.34E-01 | 9.32E-01 | -1.49E-02 | -5.46E-01 | -4.73E-01 |
| JCF_R | 1.11E+01 | 5.89E+00 | -5.00E+00 | 1.54E+00 | 1.38E+01 | -3.99E+00 | -9.30E+00 | -5.80E+00 |
| JCF_FFT | 5.95E+00 | -5.79E+01 | -2.20E+01 | 3.30E+01 | 4.99E+01 | 1.30E+01 | -2.45E+01 | -2.86E+01 |
| RXF_FE | 2.49E+02 | 2.39E+03 |  | -2.62E+03 | 6.04E+03 | 2.43E+03 | -1.58E+03 | -6.18E+01 |
| RXF_FE_FFT | -1.77E-05 | -2.64E-05 | 2.79E-05 |  | 3.32E-05 | 2.31E-05 |  | -1.85E-05 |
| RXF_SMR | -9.92E-02 | 4.87E-02 | 2.85E-02 |  | 7.53E-02 | 4.70E-02 | -1.05E-01 | -7.57E-02 |
| JCF_FE | 2.73E+03 | 5.32E+03 | 2.81E+03 | -1.44E+04 | 2.48E+04 | 8.26E+03 | -3.13E+03 | 2.08E+03 |
| JCF_FE_FFT | -4.55E-05 | -8.76E-05 | 1.63E-04 | -7.13E-05 | 1.98E-04 | 9.69E-05 |  | -9.41E-05 |
| JCF_SMR |  |  |  |  | 2.76E-01 | 4.85E-02 |  |  |

**Table A3-2.** Continued LASSO regression model outcome coefficients, bolded values are significant with p < 0.05 (Range Of Motion = ROM). Outcome metrics in lower complexity values were added as possible predictors in each model.

| Outcome | DIS | IMU_FFT | RXF | RXF_R | RXF_FFT | JCF | JCF_R | JCF_FFT |
| --- | --- | --- | --- | --- | --- | --- | --- | --- |
| RXF | 4.86E-02 |  |  |  |  |  |  |  |
| RXF_R | 7.74E+00 |  |  |  |  |  |  |  |
| RXF_FFT | 4.43E-01 | 1.48E+00 |  |  |  |  |  |  |
| JCF | -6.44E-01 |  | 9.49E-01 | -1.21E+00 | 1.48E+00 |  |  |  |
| JCF_R | -1.01E+01 |  | 3.24E+01 | -2.44E+01 | 2.38E+01 |  |  |  |
| JCF_FFT | -2.41E+01 |  | 8.20E+00 | -2.09E+01 | 5.89E+01 |  |  |  |
| RXF_FE | 4.43E+02 |  | 2.09E+04 | -2.16E+03 | 4.64E+03 | -1.08E+04 | -3.15E+03 | 7.61E+03 |
| RXF_FE_FFT |  |  | 3.66E-05 | 3.54E-05 | 1.36E-05 | -5.52E-05 | -3.07E-05 | 1.03E-04 |
| RXF_SMR | 7.06E-02 | -3.33E-02 | 3.84E-01 | 5.26E-01 | -3.50E-02 | -1.01E-01 | -1.18E-01 | 1.72E-01 |
| JCF_FE | 1.77E+03 |  | 6.65E+03 | -5.88E+03 | 5.84E+03 | 1.89E+04 | -1.05E+04 | 3.34E+04 |
| JCF_FE_FFT | 2.91E-04 | -3.15E-04 | 1.62E-04 | 1.27E-04 | -4.19E-04 | -3.42E-04 | -1.68E-04 | 1.12E-03 |
| JCF_SMR | -3.67E-02 |  |  |  |  | 1.18E-01 | 1.43E+00 | 8.40E-01 |

**Table A4.** Testing results for full LASSO regression models.

| Outcome | Mean | Standard deviation ($\sigma$) | M.S.E. | M.A.E. | M.A.E. / $\sigma$ | $r^2$ |
| --- | --- | --- | --- | --- | --- | --- |
| **RXF** | 2.87E+00 | 9.61E-01 | 3.47E-01 | 4.92E-01 | 51.2% | 0.52 |
| **RXF_R** | 1.26E+02 | 7.12E+01 | 3.20E+03 | 4.28E+01 | 60.1% | 0.24 |
| **RXF_FFT** | 1.10E+02 | 3.13E+01 | 1.77E+02 | 1.14E+01 | 36.6% | 0.79 |
| **JCF** | 1.08E+01 | 2.89E+00 | 1.20E+00 | 8.66E-01 | 29.9% | 0.75 |
| **JCF_R** | 1.66E+02 | 5.47E+01 | 5.90E+02 | 1.96E+01 | 35.8% | 0.67 |
| **JCF_FFT** | 4.79E+02 | 1.49E+02 | 3.58E+03 | 4.71E+01 | 31.6% | 0.83 |
| **RXF_FE** | 6.65E+04 | 2.49E+04 | 2.88E+07 | 4.12E+03 | 16.6% | 0.92 |
| **RXF_FE_FFT** | 4.08E-04 | 1.53E-04 | 1.88E-09 | 3.44E-05 | 22.5% | 0.88 |
| **RXF_SMR** | 9.86E-01 | 9.12E-01 | 1.85E-02 | 1.09E-01 | 12.0% | 0.94 |
| **JCF_FE** | 2.43E+05 | 6.81E+04 | 4.14E+08 | 1.52E+04 | 22.4% | 0.91 |
| **JCF_FE_FFT** | 1.74E-03 | 7.54E-04 | 4.49E-08 | 1.64E-04 | 21.7% | 0.94 |
| **JCF_SMR** | 4.33E+00 | 2.55E+00 | 2.34E-01 | 3.80E-01 | 14.9% | 0.95 |

## References

1. Bouxsein ML, Eastell R, Lui LY, et al. Change in Bone Density and Reduction in Fracture Risk: A Meta-Regression of Published Trials. Journal of Bone and Mineral Research. 2019;34(4):632–642. doi:10.1002/JBMR.3641

2. Salari N, Ghasemi H, Mohammadi L, et al. The global prevalence of osteoporosis in the world: a comprehensive systematic review and meta-analysis. J Orthop Surg Res. 2021;16(1):609. doi:10.1186/s13018-021-02772-0

3. Williams SA, Daigle SG, Weiss R, Wang Y, Arora T, Curtis JR. Economic Burden of Osteoporosis-Related Fractures in the US Medicare Population. Annals of Pharmacotherapy. 2021;55(7):821–829. doi:10.1177/1060028020970518

4. Lewiecki EM. Osteoporosis: Clinical Evaluation. (Feingold KR, Anawalt B, Blackman MR, eds.). MDText.com, Inc.; 2021.

5. Li H, Xiao Z, Quarles LD, Li W. Osteoporosis: Mechanism, Molecular Target and Current Status on Drug Development. Curr Med Chem. 2021;28(8):1489–1507. doi:10.2174/0929867327666200330142432

6. Heaney RP, Abrams S, Dawson-Hughes B, et al. Peak Bone Mass. Osteoporosis International. 2001;11(12):985–1009. doi:10.1007/s001980070020

7. Burr DB, Robling AG, Turner CH. Effects of biomechanical stress on bones in animals. Bone. 2002;30(5):781–786. doi:10.1016/S8756-3282(02)00707-X

8. Frost HM. Bone “mass” and the “mechanostat”: A proposal. Anat Rec. 1987;219(1):1–9. doi:10.1002/AR.1092190104

9. Frost HM. Bone’s mechanostat: A 2003 update. Anat Rec A Discov Mol Cell Evol Biol. 2003;275A(2):1081–1101. doi:10.1002/AR.A.10119

10. Vasto S, Amato A, Proia P, Caldarella R, Cortis C, Baldassano S. Dare to jump: The effect of the new high impact activity SuperJump on bone remodeling. A new tool to maintain fitness during COVID-19 home confinement. Biol Sport. 2021;39(4):1011–1019. doi:10.5114/BIOLSPORT.2022.108993

11. Zhao R, Zhao M, Zhang L. Efficiency of Jumping Exercise in Improving Bone Mineral Density Among Premenopausal Women: A Meta-Analysis. Published online 2014. doi:10.1007/s40279-014-0220-8

12. Srinivasan S, Weimer DA, Agans SC, Bain SD, Gross TS. Low-Magnitude Mechanical Loading Becomes Osteogenic When Rest Is Inserted Between Each Load Cycle. Journal of Bone and Mineral Research. 2002;17(9):1613–1620. doi:10.1359/jbmr.2002.17.9.1613

13. Carter DR, Orr TE. Skeletal development and bone functional adaptation. Journal of Bone and Mineral Research. 1992;7(S2):S389–S395. doi:10.1002/jbmr.5650071405

14. Ahola R, Korpelainen R, Vainionpää A, Jämsä T. Daily impact score in long-term acceleration measurements of exercise. J Biomech. 2010;43(10):1960–1964. doi:10.1016/J.JBIOMECH.2010.03.021

15. Adams DJ, Spirt AA, Brown TD, Fritton SP, Rubin CT, Brand RA. Testing the daily stress stimulus theory of bone adaptation with natural and experimentally controlled strain histories. J Biomech. 1997;30(7):671–678. doi:10.1016/S0021-9290(97)00004-3

16. Valdimarsson O, Linden C, Johnell O, Gardsell P, Karlsson MK. Daily physical education in the school curriculum in prepubertal girls during 1 year is followed by an increase in bone mineral accrual and bone width--data from the prospective controlled Malmö pediatric osteoporosis prevention study. Calcif Tissue Int. 2006;78(2):65–71. doi:10.1007/S00223-005-0096-6/METRICS

17. Turner CH. Three rules for bone adaptation to mechanical stimuli. Bone. 1998;23(5):399–407. doi:10.1016/S8756-3282(98)00118-5

18. McNamara BP, Prendergast PJ, Taylor D. Prediction of bone adaptation in the ulnar-osteotomized sheep’s forelimb using an anatomical finite element model. J Biomed Eng. 1992;14(3):209–216. doi:10.1016/0141-5425(92)90054-O

19. Webster D, Schulte FA, Lambers FM, Kuhn G, Müller R. Strain energy density gradients in bone marrow predict osteoblast and osteoclast activity: A finite element study. J Biomech. 2015;48(5):866–874. doi:10.1016/j.jbiomech.2014.12.009

20. MacNeil JA, Boyd SK. Bone strength at the distal radius can be estimated from high-resolution peripheral quantitative computed tomography and the finite element method. Bone. 2008;42(6):1203–1213. doi:10.1016/J.BONE.2008.01.017

21. Troy KL, Mancuso ME, Johnson JE, Wu Z, Schnitzer TJ, Butler TA. Bone Adaptation in Adult Women Is Related to Loading Dose: A 12-Month Randomized Controlled Trial. J Bone Miner Res. 2020;35(7):1300–1312. doi:10.1002/JBMR.3999

22. Mancuso ME, Wilzman AR, Murdock KE, Troy KL. Effect of external mechanical stimuli on human bone: a narrative review. Progress in Biomedical Engineering. 2022;4(1):012006. doi:10.1088/2516-1091/ac41bc

23. Hsieh YF, Wang T, Turner CH. Viscoelastic response of the rat loading model: implications for studies of strain-adaptive bone formation. Bone. 1999;25(3):379–382. doi:10.1016/S8756-3282(99)00181-7

24. Seth A, Hicks JL, Uchida TK, et al. OpenSim: Simulating musculoskeletal dynamics and neuromuscular control to study human and animal movement. PLoS Comput Biol. 2018;14(7):e1006223. doi:10.1371/journal.pcbi.1006223

25. Delp SL, Anderson FC, Arnold AS, et al. OpenSim: Open-Source Software to Create and Analyze Dynamic Simulations of Movement. IEEE Trans Biomed Eng. 2007;54(11):1940–1950. doi:10.1109/TBME.2007.901024

26. ANDERSON FC, PANDY MG. A Dynamic Optimization Solution for Vertical Jumping in Three Dimensions. Comput Methods Biomech Biomed Engin. 1999;2(3):201–231. doi:10.1080/10255849908907988

27. Kawalilak CE, Kontulainen SA, Amini MA, Lanovaz JL, Olszynski WP, Johnston JD. In vivo precision of three HR-pQCT-derived finite element models of the distal radius and tibia in postmenopausal women. BMC Musculoskelet Disord. 2016;17(1):389. doi:10.1186/s12891-016-1238-x

28. Weinhandl JT, Joshi M, O’Connor KM. Gender Comparisons between Unilateral and Bilateral Landings. J Appl Biomech. 2010;26(4):444–453. doi:10.1123/jab.26.4.444

29. Sara LK, Gaudette LW, Souza Júnior JR de, Tenforde AS, Wasserman L, Johnson CD. Cues to land softly and quietly result in acute reductions in ground reaction force loading rates in runners. Gait Posture. 2024;109:220–225. doi:10.1016/j.gaitpost.2024.02.008

30. Vlachopoulos D, Barker AR, Ubago-Guisado E, Williams CA, Gracia-Marco L. The effect of a high-impact jumping intervention on bone mass, bone stiffness and fitness parameters in adolescent athletes. Arch Osteoporos. 2018;13(1):128. doi:10.1007/s11657-018-0543-4

31. Zhao R, Zhao M, Zhang L. Efficiency of Jumping Exercise in Improving Bone Mineral Density Among Premenopausal Women: A Meta-Analysis. Sports Medicine. 2014;44(10):1393–1402. doi:cvb

32. Nguyen VH. School-based exercise interventions effectively increase bone mineralization in children and adolescents. Osteoporos Sarcopenia. 2018;4(2):39–46. doi:10.1016/j.afos.2018.05.002

33. Burt LA, Liang Z, Sajobi TT, Hanley DA, Boyd SK. Sex- and Site-Specific Normative Data Curves for HR-pQCT. Journal of Bone and Mineral Research. 2016;31(11):2041–2047. doi:10.1002/jbmr.2873

34. Szulc P, Dufour AB, Hannan MT, et al. Fracture risk based on high-resolution peripheral quantitative computed tomography measures does not vary with age in older adults—the bone microarchitecture international consortium prospective cohort study. Journal of Bone and Mineral Research. Published online March 3, 2024. doi:10.1093/jbmr/zjae033

35. Roelker SA, Caruthers EJ, Hall RK, Pelz NC, Chaudhari AMW, Siston RA. Effects of Optimization Technique on Simulated Muscle Activations and Forces. J Appl Biomech. 2020;36(4):259–278. doi:10.1123/jab.2018-0332

36. MacNeil JA, Boyd SK. Bone strength at the distal radius can be estimated from high-resolution peripheral quantitative computed tomography and the finite element method. Bone. 2008;42(6):1203–1213. doi:10.1016/j.bone.2008.01.017

37. Johnson JE, Troy KL. Validation of a new multiscale finite element analysis approach at the distal radius. Med Eng Phys. 2017;44:16–24. doi:10.1016/j.medengphy.2017.03.005

